# Membrane structural properties in *Staphylococcus aureus* are tuned by the carotenoid 4,4′-diaponeurosporenoic acid

**DOI:** 10.64898/2026.04.08.716698

**Authors:** Jessica Múnera-Jaramillo, Gerson-Dirceu López, Elizabeth Suesca, Elena Ibáñez, Alejandro Cifuentes, Chiara Carazzone, Chad Leidy, Marcela Manrique-Moreno

## Abstract

*Staphylococcus aureus* (*S. aureus*) is a clinically relevant pathogen capable of adapting its membrane composition in response to environmental stress. In this adaptive process, bacterial carotenoids play a crucial role. Although staphyloxanthin (STX) is the main carotenoid produced by the bacterium, *S. aureus* also synthesizes other pigmented intermediates that play an unknown role in regulating membrane biophysical properties. In this study, we purified 4,4’-diaponeurosporenoic acid (4,4′-DNPA) from *S. aureus* carotenoid extracts and evaluated its effect on the thermotropic and biophysical properties of representative membrane models. The highly rigid triterpenoid 4,4′-DNPA is one of the last precursors in the biosynthesis of STX and is found in high concentrations in the stationary phase of *S. aureus*. Phase transition temperatures were determined using infrared spectroscopy, while interfacial hydration and hydrophobic core dynamics were investigated using fluorescence spectroscopy through Laurdan generalized polarization and DPH anisotropy. The results show that 4,4′-DNPA increases the main phase transition temperature of lipid bilayers in a concentration-dependent manner. This is in contrast to STX that decreases the transition temperature. This difference is consistent with the additional fatty acid present in STX that changes its effect on the phase behavior. Furthermore, 4,4′-DNPA reduced the interfacial hydration levels and restricted hydrophobic-core dynamics at higher concentrations, consistent with increased molecular order and stability. 4,4′-DNPA therefore complements STX in increasing membrane order and lipid packing. These findings support the notion that the production of bacterial carotenoids functions as a biophysical regulatory mechanism of lipid packing in *S. aureus* membranes.

## 1. Introduction

*Staphylococcus aureus* (*S. aureus*) is an opportunistic pathogen responsible for infections ranging from mild cutaneous infections to life-threatening diseases, including pneumonia, endocarditis, and sepsis [1, 2]. Its clinical relevance has increased in recent decades due to the emergence of multidrug-resistant strains, which represents a serious global public health challenge [3, 4]. The remarkable ability of *S. aureus* to adapt to different environmental conditions is a key factor in its success as a pathogen [5, 6]. Among its adaptive strategies, *S. aureus* synthesizes a distinctive class of carotenoid pigments, primarily staphyloxanthin (STX), a C30 carotenoid with a hydrocarbon chain conjugated to a sugar and a fatty acid moiety [7–9]. These carotenoids are responsible for the characteristic golden coloration of the bacterium [7, 8, 10]. Beyond their pigmentary role, these molecules play a crucial function in the pathogenicity of *S. aureus*, contributing to resistance against oxidative stress and enhancing survival within the host [11–13]. Several studies have shown that carotenoid-deficient mutants are more susceptible to reactive oxygen species and exhibit reduced virulence, evidencing their role in protecting bacterial membranes and cellular components from oxidative damage [14–16].

In addition to their antioxidant function, carotenoids have been suggested to function as regulators of the biophysical state of *S. aureus* cell membranes [15, 17]. These molecules, due to the structural rigidity conferred by their linear, conjugated polyene chains and diverse polar headgroups, can intercalate between lipid molecules, influencing both the packing of the acyl chains and the hydration of the polar interface [18–20]. Fluorescence spectroscopy studies evaluating DPH anisotropy and Laurdan generalized polarization have shown that carotenoids induce increased order in the hydrocarbon core and reduced interfacial hydration, respectively [17, 21, 22]. These findings reflect the ability of carotenoids to modulate the acyl chain order and water penetration into the bilayer interface. Such structural modifications are thought to contribute to enhanced resistance to antimicrobial peptides (AMPs), which often exploit membrane defects or increased fluidity to permeabilize bacterial membranes [23, 24]. Furthermore, thermodynamic analysis, such as differential scanning calorimetry (DSC) experiments, have demonstrated that carotenoid-containing membranes exhibit broader and shifted phase transitions compared to carotenoid-free systems [25, 26]. These changes suggest altered cooperativity within the lipid systems and increased stability of the bilayer structure, potentially providing a more gradual and controlled response to environmental stresses [27]. These findings indicate that the biosynthesis of carotenoids in *S. aureus* not only enhances its resistance to oxidative stress but also modulates the physicochemical properties of its membranes, which may lead to increased resistance to host defense mechanisms.

Although STX is the predominant carotenoid, *S. aureus* synthesizes a broader diversity of carotenoids, including biosynthetic intermediates, whose relative abundance can vary depending on environmental stimuli [8]. This molecular diversity suggests that additional carotenoid species, apart from STX, may actively participate in modulating the biophysical properties of the membrane. In a previous work [25], we have evaluated the influence of total carotenoids produced by *S. aureus* on the thermotropic behavior of synthetic membrane systems. First, we analyzed the effect of total carotenoid extracts on the phase behavior of lipid models through a detailed thermodynamic study using Fourier-transform infrared spectroscopy (FT-IR) and differential scanning calorimetry (DSC), which demonstrated that total carotenoids regulate the biophysical properties of membranes [25]. Subsequently, we focused specifically on the main carotenoid, STX [25], and investigated its effect on the thermotropic and biophysical properties of *S. aureus* membrane models through FT-IR and fluorescence spectroscopy. These studies revealed that STX decreases the phase transition temperature and increases membrane fluidity, suggesting a modulatory role in maintaining membrane dynamics under varying conditions. Notably, during the purification and characterization of STX, a biosynthetic precursor known as 4,4′-diaponeurosporenic acid (4,4′-DNPA), was identified in stationary-phase cultures. This intermediate is formed in the STX biosynthetic pathway through the oxidation of the terminal methyl group of 4,4′-diaponeurosporene, a reaction catalyzed by the mixed-function oxidase CrtP (Fig. 1). The detection of 4,4′-DNPA as a persistent species in the stationary phase suggests that it is not merely a transient precursor but may contribute to a regulatory mechanism modulating the biophysical state of the *S. aureus* cell membrane.

**Fig. 1.**
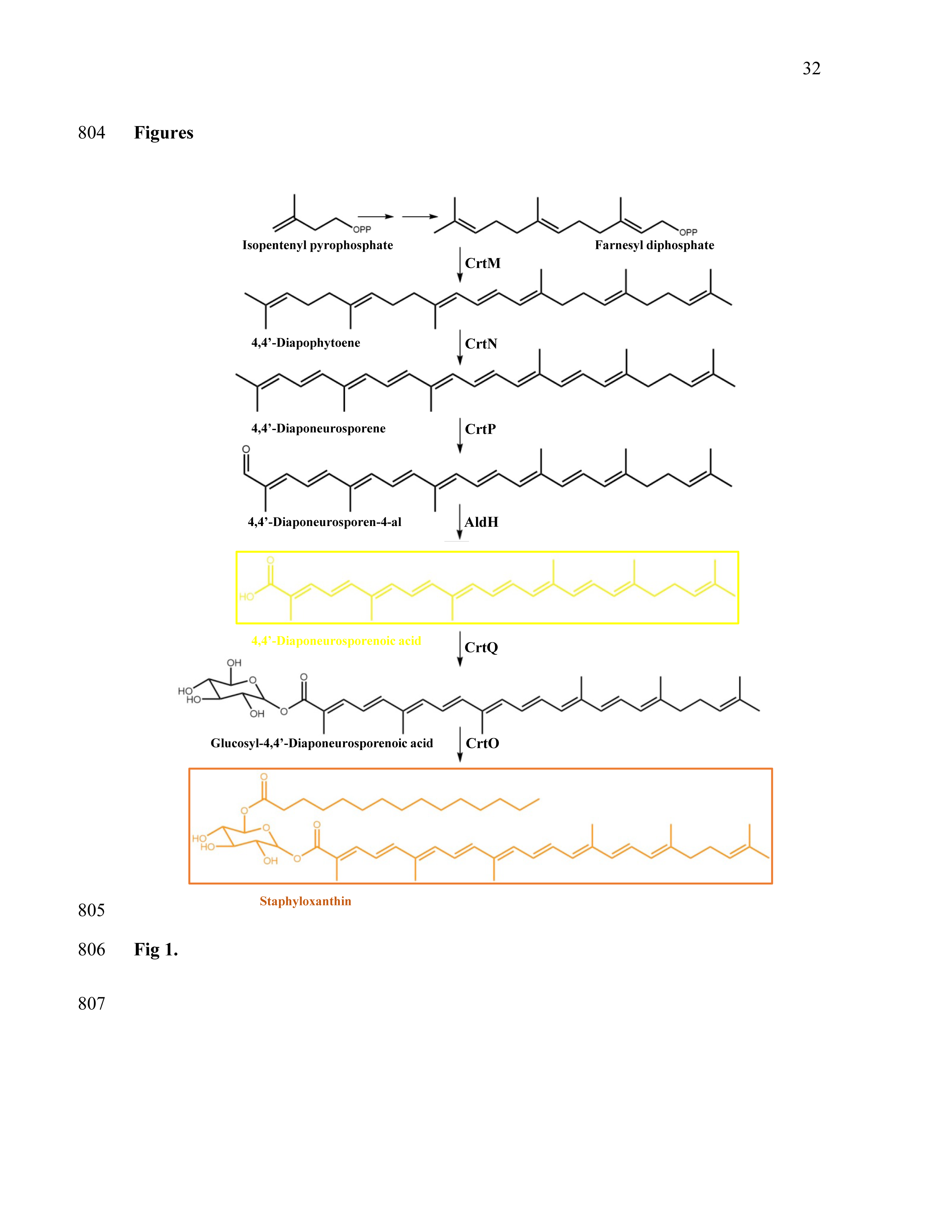
Biosynthesis pathway of STX. The biosynthesis of STX, the pathway begins with the head-to-head condensation of two molecules of farnesyl diphosphate to produce dehydrosqualene (4,4′-diapophytoene), catalyzed by the dehydrosqualene synthase (CrtM). Next, dehydrosqualene is desaturated by the enzyme dehydrosqualene desaturase (CrtN) to produce the yellow intermediate 4,4′-diaponeurosporene. Subsequently, the terminal methyl group of 4,4′-diaponeurosporene is oxidized by the mixed-function oxidase (CrtP), forming 4,4′-diaponeurosporenic acid. This acid is then glycosylated at the C1 position by the glycosyltransferase (CrtQ), producing glycosyl 4,4′-diaponeurosporenoate. Finally, the glucose moiety at the C6 position is esterified with 12-methyltetradecanoic acid by the acyltransferase (CrtO), yielding the final product, STX. The chemical structures were generated with ChemDraw 19.0 [7, 64].

To evaluate its possible role in modulating membrane biophysical properties, we purified 4,4′-DNPA from carotenoid extracts of *S. aureus* and analyzed its effects on representative lipid model systems composed of an 80:20 mixture of DMPG:CL, mimicking the main phospholipids of *S. aureus* membranes. The analysis was performed using FT-IR to determine the phase transition temperature and fluorescence spectroscopy to evaluate interfacial hydration and hydrophobic core dynamics using Laurdan Generalized Polarization and DPH anisotropy, respectively. This experimental approach offers insights into the distinct contributions of carotenoid intermediates to membrane regulation, highlighting the strategies employed by *S. aureus* to maintain its membrane integrity.

## 2. Material and Methods

### 2.1. Reagents

1,2-dimyristoyl-sn-glycero-3-phosphoglycerol sodium salt (DMPG, Lot. 140PG-167) and 1’,3’-bis[1,2-dimyristoleoyl-sn-glycero-3-phospho]-glycerol sodium salt (CL, Lot. 50332P-200MG-A-030) were purchased from Avanti Polar Lipids (Alabaster, AL, USA). 4-(2-hydroxyethyl)-1-piperazineethanesulfonic acid (HEPES) was purchased from Sigma-Aldrich (St. Louis, MO, USA), HPLC-grade methanol, ethyl acetate, and chloroform were purchased from Honeywell (Detroit, MI, USA) and J.T. Bajer (Palo Alto, CA, USA) respectively. Sodium chloride (NaCl) Reagent Plus (>99%) and butylated hydroxytoluene (BHT) were purchased from Sigma-Aldrich (St. Louis, MO, USA). Ethylenediaminetetraacetic acid (EDTA) was purchased from Amresco (Solon, OH, USA). Luria–Bertani (LB) medium was prepared with NaCl (ACS, J.T. Baker, USA), tryptone (OXOIO, Basigstoke, Hampshire, UK), and a yeast extract (Dibico, Mexico D.F., Mexico). HPLC-water was obtained from a water purification system: Heal Force Smart-Mini (Shangai, China). The fluorescent probes 6-dodecanoyl-2-dimethylaminonaphthalene (Laurdan) and diphenylhexariene (DPH) were purchased from ThermoFisher Scientific (Waltham, MA, USA).

### 2.2. Bacterial cultures

A methicillin-susceptible *S. aureus* strain (SA401) was used for bacterial cultures. An entire characterization of its carotenoid composition and biophysical characteristics was previously published [8, 21]. A single colony of the *S. aureus* strain was grown in 10 ml of LB medium at 37 °C overnight (16 h) with continual agitation (250 rpm). LB medium contained, per liter, 10 g of NaCl, 10 g of tryptone, and 5 g of yeast extract. Then, the cells were diluted (1:1000) in fresh LB medium and cultivated for 24 h. Finally, the cell pellet was obtained by centrifugation at 7300× *g* at 4 °C for 10 min (Thermo Scientific, Waltham, MA, USA) and lyophilized for 24 h (LABCONCO, Kansas City, MO, USA).

### 2.3. Carotenoid extraction

Total carotenoid extract was obtained from *S. aureus* cells using a previously reported conventional extraction method [8, 29]. In addition, pressurized liquid extraction (PLE) was evaluated using an accelerated solvent extractor (ASE 200, Dionex, Sunnyvale, CA, USA).For PLE, 1 g of lyophilized and ground *S. aureus* cells was loaded into an 11 ml extraction cell and extracted with ethanol in static mode for 20 min at 100 bars and a temperature of 125 °C. The solvent was evaporated under a gentle stream of nitrogen at room temperature, and the extracts were stored at −20 °C until further processing. Carotenoid extracts obtained by both methods (conventional and PLE) were subsequently resuspended in methanol and subjected to liquid-liquid extraction by adding ethyl acetate and a 1.7 M solution of NaCl (1:3 v/v). The mixture was vortexed and centrifugated at 7300 × *g* for 15 min at 4 °C to promote phase separation. The organic phase was collected and concentrated using a rotary evaporator in a water bath at 35 °C. To further remove non-lipid contaminants, the extract was resuspended in 20 mL of chloroform:methanol (2:1, v/v) and transferred to a borosilicate glass tube with a phenolic cap and polytetrafluoroethylene (PTFE)-faced liner. Subsequently, 7 mL of 1.7 M NaCl solution was added, the mixture was shaken, and phase separation was induced by centrifugation at 630 × *g* for 15 min at 4 °C. The lower chloroform phase containing carotenoids was collected, dried over anhydrous sodium sulfate, transferred to amber tubes, and evaporated to dryness using a refrigerated vacuum concentrator (CentriVap, LAB-CONCO, Kansas City, MO, USA).

### 2.4. 4,4’-DNPA purification and identification

Purification of 4,4’-DNPA was carried out following a previously reported method [28]. Briefly, the carotenoid extract was applied onto preparative thin-layer chromatography (PTLC) plates (10 x20 cm), and separations were performed using two mobile phases: toluene:methanol:acetic acid (87:11:2; v/v) and toluene:methanol (84:16). After development (25–30 min), the band corresponding to 4,4′-DNPA was recovered from the silica using methanol:chloroform (2:1, v/v), dried over anhydrous sodium sulfate, and concentrated to dryness using a refrigerated vacuum concentrator.

Putative identification of 4,4′-DNPA was performed using a previously reported LC-MS method [8]. The dried extract was reconstituted at 10 mg mL⁻¹ in chloroform:methanol (1:2, v/v) and analyzed using a Dionex UltiMate 3000 UHPLC system equipped with a diode-array detector (DAD) and coupled to an LCQ Fleet ion trap mass spectrometer via an atmospheric pressure chemical ionization (APCI) source operated in negative ion mode. Data acquisition and processing were performed using Xcalibur 4.3 software (Thermo Scientific, San Jose, CA, USA). Chromatographic separation was carried out on a YMC-C30 column (150 × 4.6 mm, 3 μm; YMC America, Inc., Devens, MA, USA) protected with a C18 guard cartridge (4 × 2 mm, 3 μm; Phenomenex). A 10 μL injection volume was used, with samples maintained at 5 °C in the autosampler. The mobile phases consisted of methanol:methyl tert-butyl ether:water (80:18:2, v/v/v; solvent A) and methanol:methyl tert-butyl ether:water (8:89:3, v/v/v; solvent B), both containing 400 mg L⁻¹ ammonium acetate. The flow rate was 0.45 mL min⁻¹. The gradient elution was as follows: 5% B (0–3 min), 5–10% B (3–9 min), 10–25% B (9–19 min), 25–40% B (19–23 min), held at 40% B (23–27 min), increased to 100% B (27–31 min), held at 100% B (31–34 min), returned to 5% B (34–36 min), and equilibrated at 5% B until 40 min. DAD data were acquired over the UV–Vis range (240–600 nm), and carotenoid absorbance was monitored between 400 and 500 nm. The APCI source was operated under the following conditions: vaporizer temperature 300 °C; discharge current 15.0 μA; capillary voltage −25.0 V; tube lens −80.0 V; capillary temperature 350 °C; sheath gas flow 50 arbitrary units; and auxiliary gas flow 30 arbitrary units. The ion trap was operated in full-scan mode (m/z 65–1200) and data-dependent MS/MS mode using 30% collision energy and an isolation width of 3 m/z. The assignment of 4,4′-DNPA was based on its chromatographic behavior, characteristic UV–Vis absorption profile, and APCI-MS/MS fragmentation pattern, in agreement with previously reported data [28].

### 2.5. Infrared spectroscopy experiments

Supported lipid bilayers (SLBs) were prepared *in situ* in a BioATR II cell. The unit was integrated with a Tensor II spectrometer (Bruker Optics, Ettlingen, Germany) with a liquid nitrogen MCT detector using a spectral resolution of 4 cm^-1^ and 120 scans per spectrum. The desired temperature was set by a Huber Ministat 125 computer-controlled circulating water bath (Huber, Offenburg, Germany) with an accuracy of ± 0.1 °C. The background was taken using HEPES buffer 20 mM, 500 mM NaCl, and 1 mM EDTA in the same temperature range as the tested lipid systems. For *in situ* measurements, stock solutions of pure DMPG, CL, and DMPG:CL (80:20) were prepared in chloroform. Subsequently, the cell was filled with 20 µl of a 20 mM lipid stock solution, and 4,4’-DNPA was added in different concentrations (5, 10, 15, and 20 mol%); then, the chloroform was evaporated, resulting in a lipid film. Afterward, the cell was filled with 20 µL buffer solution and incubated over the phase transition temperature for 10 min. To determine the position of the vibrational band in the range of the second derivate of the spectra, all the absorbance spectra were cut in the 2970-2820 cm^-1^ range, shifted to a zero baseline, and the peak-picking function included in OPUS 7.5 software (Bruker Optics, Ettlingen, Germany) was used. The results were plotted as a function of the temperature. Finally, to determine the transition temperature (Tm) of the lipids, the curve was fitted according to the Boltzmann model to calculate the inflection point of the obtained thermal transition curves using the OriginPro 8.0 software (OriginLab Corporation, Northampton, MA, USA).

### 2.6. Fluorescence spectroscopy experiments

To perform fluorescence measurements, DMPG:CL (80:20) multilamellar vesicles (MLVs) were dissolved in chloroform containing 5% v/v methanol with varying concentrations of 4’4’-DNPA incorporated (molar percentages of 5, 10, 15, and 20%); subsequently DPH and Laurdan dyes were added at a concentration of 1:150 (probe to lipid ratio). The mixtures were dried under a stream of nitrogen and lyophilized overnight to remove residual chloroform [21]. Then, the obtained films were resuspended using 500 µl of a 20 mM HEPES, 400 mOsm buffer at 50 °C to a final concentration of 3 mM under periodic agitation using a vortex mixer. For all measurements, 30 µl of MLVs containing the fluorescent probes were added to a cuvette containing 1 ml of HEPES buffer.

Laurdan generalized polarization (Laurdan GP) and DPH anisotropy (r) measurements were acquired in an ISS-PC1 (ISS, Champaign, IL, USA) photon-counting spectrofluorometer equipped with a temperature controller and continuous magnetic stirring [21]. A combination of 0.5-2.0 mm slits was used in the excitation and emission monochromators to set the bandpass to 2, 4, or 8 nm, depending on the sample intensity and photobleaching sensitivity [21]. The Laurdan GP value was calculated using the equation 1:

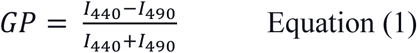

Where *I_440_* and *I_490_* are the fluorescence intensities at emission wavelengths of 440 nm (solid-ordered phase) and 490 nm (liquid-disordered phase), respectively [29].

On the other hand, DPH fluorescence anisotropy was calculated according to the equation 2:

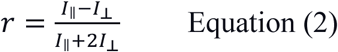

Where *I*_ǁ_is the fluorescence intensity when the angle between polarizers is 0° and *I*_&_ is the fluorescence intensity when the angle between polarizers is 90° [29]. Fluorescence measurements were performed from 10 to 45 °C (heat rate: 0.5 °C/min). The data presented in the figures represent mean values and standard error of 4 measurements in four independent experiments.

## 3. Results

### 3.1. 4,4’-DNPA purification and identification

To isolate and characterize the biosynthetic precursor 4,4’-DNPA, total carotenoid extracts were obtained from *S. aureus* cells harvested at the stationary phase of growth. The extraction method previously reported by our group [8, 29], which efficiently recovers bacterial carotenoids, was employed in this study. Purification of 4,4’-DNPA was achieved by preparative thin-layer chromatography (PTLC), which enabled separation of the biosynthetic precursor (R_F_ = 0.48) and staphyloxanthin (R_F_ = 0.29), indicating the coexistence of both compounds at late stages of the biosynthetic pathway [8]. Representative PTLC results supporting the isolation of 4,4’-DNPA are shown in the supplementary material (Fig. S1). Putative identification of 4,4’-DNPA was performed by LC-DAD-APCI-MS/MS (Fig. 2). According to previous studies, STX is the main product of the carotenoid biosynthetic pathway in *S. aureus* [7, 8]. However, the chromatographic profile of the total carotenoid extract (Fig. 2A) reveals the presence of multiple chemical species, indicating the coexistence of STX with other carotenoids and biosynthetic intermediates. This observation is consistent with previous reports describing the diversity of carotenoid species produced by *S. aureus* under different growth conditions [8]. Specifically, the chromatogram in Fig. 2A shows two notable signals at 10.55 and 15.71, which are attributed to STX (based on LC-DAD-APCI-MS/MS data, see Fig. S2) and 4,4’-DNPA, respectively. These findings are consistent with studies indicating that *S. aureus* produces a variety of metabolites during STX biosynthesis, including STX-homologs, dehydro-STX, dehydro-STX-homologs, and other pigmented intermediates [8]. The chromatogram obtained after the purification process (Fig. 2B) shows a predominant signal at 10.63 min, indicating enrichment of 4,4’-DNPA in the purified fraction. The assignment of 4,4′-DNPA was further supported by APCI-MS data (Fig. 2C), which showed a dominant ion at *m/z* 432 corresponding to the deprotonated molecule [M–H]⁻, in agreement with previous reports [8].

**Fig. 2.**
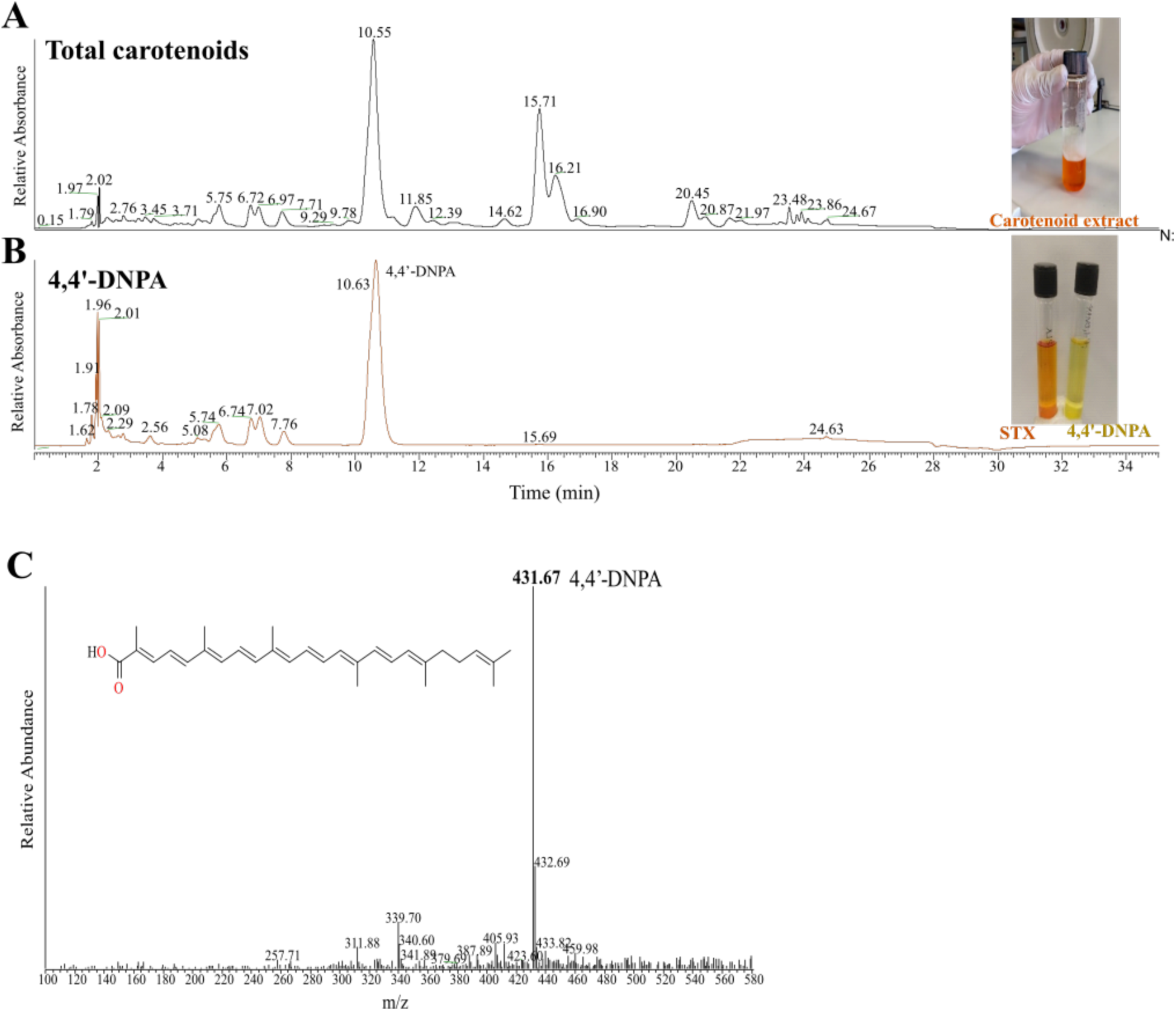
Characterization of 4,4’-DNPA by liquid chromatography coupled with mass spectrometry. **(A)** HPLC-MS analysis for a total carotenoid extracted from *S. aureus cells*, **(B)** HPLC-MS for purified 4,4’-DNPA. **(C)** Full MS spectra of 4,4’-DNPA. Chromatograms (A) and (B) were obtained at 460 nm.

### 3.2. Phase transition measurements by infrared spectroscopy

Infrared spectroscopy is used to determine the P_ý’_ gel (solid-ordered) to the L_α_ liquid-crystalline (liquid-disordered) phase transition temperature. The FT-IR technique enables monitoring the methylene symmetric stretching vibrational mode (ν_’_CH_2_), which reflects changes in the conformational order as a lipid system transitions from a highly ordered phase (solid-ordered phase, P_ý’_) to a fluid phase (liquid-disordered phase, L_α_) with increasing temperature [30, 31]. Native *S. aureus* membranes present cooperative melting phase transitions around 15°C [8, 32, 33]. This phase transition is sensitive to carotenoid content [8] and its temperature can shift based on oxygen content during growth conditions [33] and other environmental conditions that affects carotenoid levels. We evaluated the effect of several 4,4’-DNPA concentrations in the phase transition temperature of both individual phospholipid systems (DMPG and CL) and the representative lipid model of *S. aureus* cell membrane, which is built by DMPG:CL (80:20). The synthetic lipids DMPG and CL were chosen to reflect the saturated phospholipids synthesized by the bacterium [21]. Changes in wavenumber values as a function of temperature are presented in Fig 3. For the pure DMPG system, 4,4′-DNPA at 5% and 10% slightly decreased the T_m_ (Fig. 3A). In contrast, the highest concentration tested (20% 4,4’-DNPA) increased the temperature by at least 6.3 °C (Table 1), suggesting a rigidification of the DMPG lipid bilayers. Comparison of the DMPG control system with the 20% 4,4′-DNPA system reveals notable differences: whereas the phase transition in the control model is characterized by a highly cooperative state change, the pigmented model exhibits a more gradual transition (Fig. 3A). On the other hand, the DMPG systems exhibit an increase in the acyl chain order both in the solid-ordered phase regime below the T_m_, and in the liquid-disordered phase regime above T_m_ (liquid-disordered phase) as a function of the pigmented precursor concentration.

**Fig. 3.**
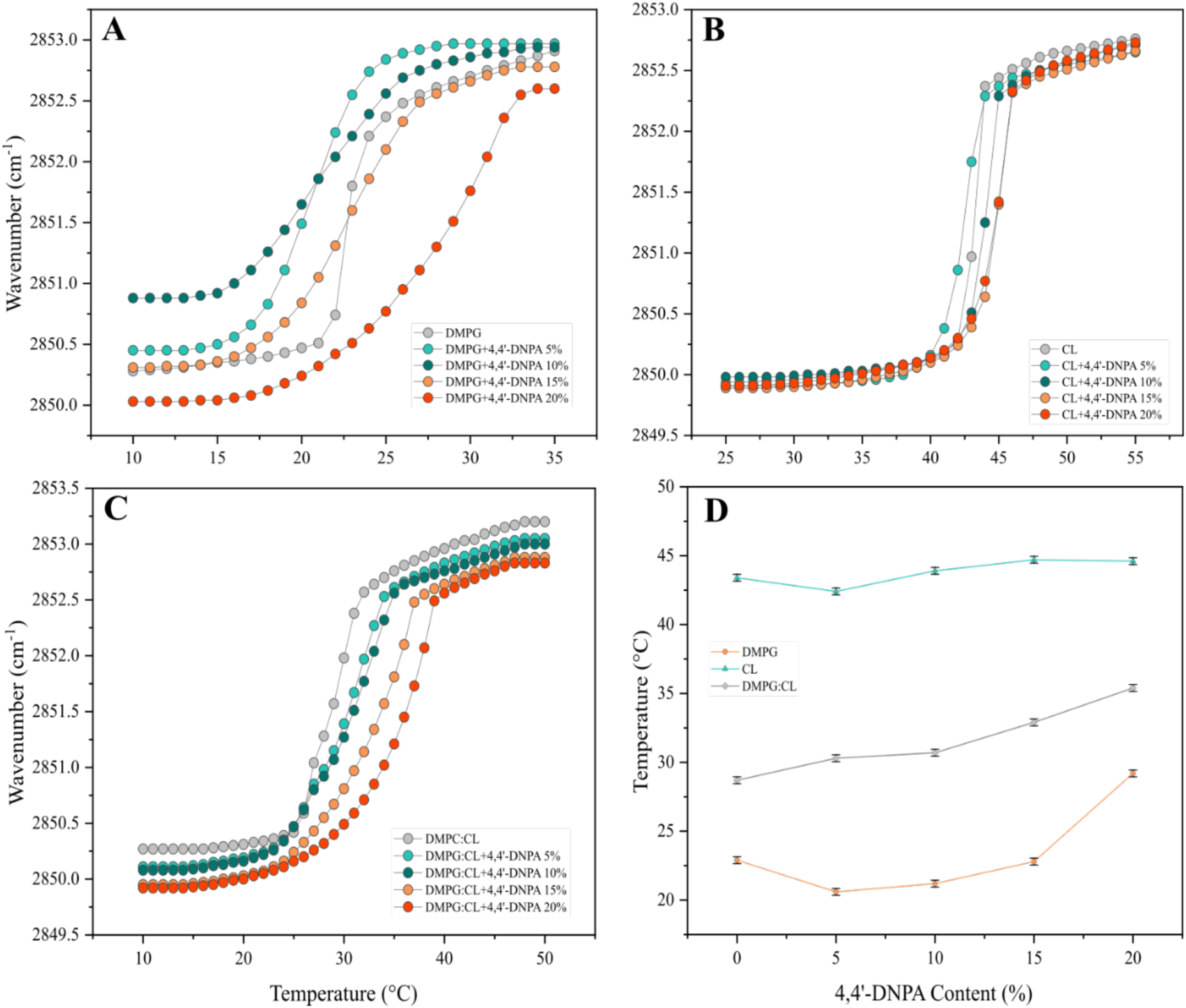
Peak positions of the νCH_2_ symmetric stretching vibrations band of the acyl chain methylene groups as a function of temperature in the presence of different concentrations of 4,4’-DNPA for **(A)** DMPG, **(B)** CL, **(c)** DMPG:CL (80:20) models, and **(D)** Temperature variation of the main phase transition as a function of 4,4’-DNPA content in DMPG, CL, and DMPG:CL (80:20) systems.

**Table 1.** Phase transition (T_m_) temperatures, of the supported bilayers of DMPG, CL, and DMPG:CL (80:20) by FTIR. Standard deviations are ≤ 0.1 °C.

| 4,4'-DNPA (% mol)/Lipid System | $T_m$ (°C) | | |
| --- | --- | --- | --- |
|  | DMPG | CL | DMPG:CL |
| 0 | 22.9 | 43.4 | 28.7 |
| 5 | 20.6 | 42.4 | 30.3 |
| 10 | 21.2 | 43.9 | 30.7 |
| 15 | 22.8 | 44.7 | 32.9 |
| 20 | 29.2 | 44.6 | 35.4 |

**Table 2.** Phase transition (T_m_) temperatures, of the supported bilayers of representative model systems including STX and 4,4’-DNPA by FTIR. Standard deviations are ≤ 0.1 °C.

| Systems | $T_m$ (°C) |
| --- | --- |
| DMPG:CL | 28.7 |
| DMPG:CL+STX/4,4'-DNPA 0/100 | 35.4 |
| DMPG:CL+STX/4,4'-DNPA 75/25 | 29.5 |
| DMPG:CL+STX/4,4'-DNPA 85/15 | 28.6 |
| DMPG:CL+STX/4,4'-DNPA 100/0 | 27.0 |

In pure CL models, a slight decrease in T_m_ value is observed at 5% 4,4’-DNPA; at the same time, higher pigment concentrations (10 to 20 molar% of 4,4’-DNPA) lead to a modest increase in the phase transition temperature compared to the control system (Table 1). Moreover, significant changes in wavenumber values at fixed temperatures are not observed (Fig. 3B). The above is consistent with previous reports indicating that lipid systems that include CL are less susceptible to mechanical perturbation by the action of external molecules or other membrane components [34, 35].

In the representative model systems of the *S. aureus* lipid bilayer, DMPG:CL (80:20), the effect of increasing 4,4’-DNPA concentrations on the wavenumber values are shown in Fig. 3C. These systems were constructed based on the previously reported lipid composition of the cell membranes of *S. aureus* [34]. The models exhibit a progressive increase in T_m_ values as the concentration of bacterial pigment increases, where the most pronounced effect is observed in the 20 mol% 4,4’-DNPA system, in which T_m_ is elevated by 6.7 °C compared to the control system (Table 1). Notably, despite containing both DMPG and CL phospholipids, the magnitude of the effect of the pigment is comparable to that observed in the pure DMPG models. In addition, these results demonstrate that the inclusion of the pigment increases the rigidity of the membrane models in a concentration-dependent manner. Furthermore, at fixed temperatures above and below the phase transition, 4,4′-DNPA increases the wavenumber values, suggesting that higher pigment concentrations enhance acyl chain order (Fig. 3c).

The data show that increasing 4,4′-DNPA content differentially affects the phase transition temperature of the lipid systems. CL-rich membranes exhibit consistently high transition temperatures with only minor fluctuations, indicating relative insensitivity to DNPA. In contrast, DMPG displays a non-linear response, with an initial decrease followed by a marked increase at higher DNPA levels. The DMPG:CL mixture shows a steady, progressive rise in transition temperature as DNPA increases. Overall, these results suggest that 4,4′-DNPA enhances membrane order in mixed systems while exerting lipid-specific effects depending on composition (Fig. 3D).

Finally, the phase transition temperature was evaluated in lipid systems composed of representative phospholipids DMPG and CL, the biosynthetic precursor 4,4’-DNPA, and STX, the main carotenoid produced by *S. aureus*. This experiment was designed to evaluate how the equilibrium ratio between 4,4’-DNPA and STX could modulate biophysical properties of the membrane. The systems were prepared with a ratio of DMPG:CL (80:20) and a carotenoid concentration of 20% concerning the total lipid composition. Additionally, the systems evaluated considered the proportions between carotenoids of 75/25 and 85/15 for STX and 4,4’-DNPA, respectively. The previously described is reasonable considering the reports indicating that *S. aureus* pigmented strains produce up to 20% of total carotenoid content [17, 25].

Thermotropic infrared spectroscopy measurements of multicomponent mixtures of DMPG:CL:STX:4,4′-DNPA are presented in Fig. 4A. The inclusion of STX in 4,4′-DNPA-containing systems decreased the phase transition temperature, indicating that STX facilitates the lipid phase change at lower temperatures compared to systems containing only 4,4′-DNPA, likely by destabilizing the solid-ordered phase. Fig. 4B plots the shifts in T_m_ of the DMPG:CL mixture in the presence of either STX or 4,4′-DNPA, showing that each molecule has an opposite effect on the T_m_. For the combined system DMPG:CL:STX:4,4′-DNPA, which includes both carotenoids, we observe a close to linear relation of the T_m_ as a function of the proportion of STX:4,4′-DNPA present in the mixture. The intermediate proportions of STX:4,4′-DNPA that were chosen reflect the proportions of these two carotenoids found in cell membranes of *S. aureus* [8]. These findings demonstrate that the combined presence of both pigments modulates the phase transition temperature in a manner that is proportional to the ratio of the two molecules present in the membrane (Fig. 4C). Moreover, at fixed temperatures, the wavenumber values remain nearly constant for all lipid systems for compositions that reflect the proportions found in the membrane (Fig. 4A). It is important to point out that, overall, the data confirms that staphyloxanthin has a strong tendency to depresses the phase transition temperature even in the presence of its biosynthetic precursor 4,4′-DNPA, even if this precursor has the tendency to increase the phase transition by itself. [28].

**Fig. 4.**
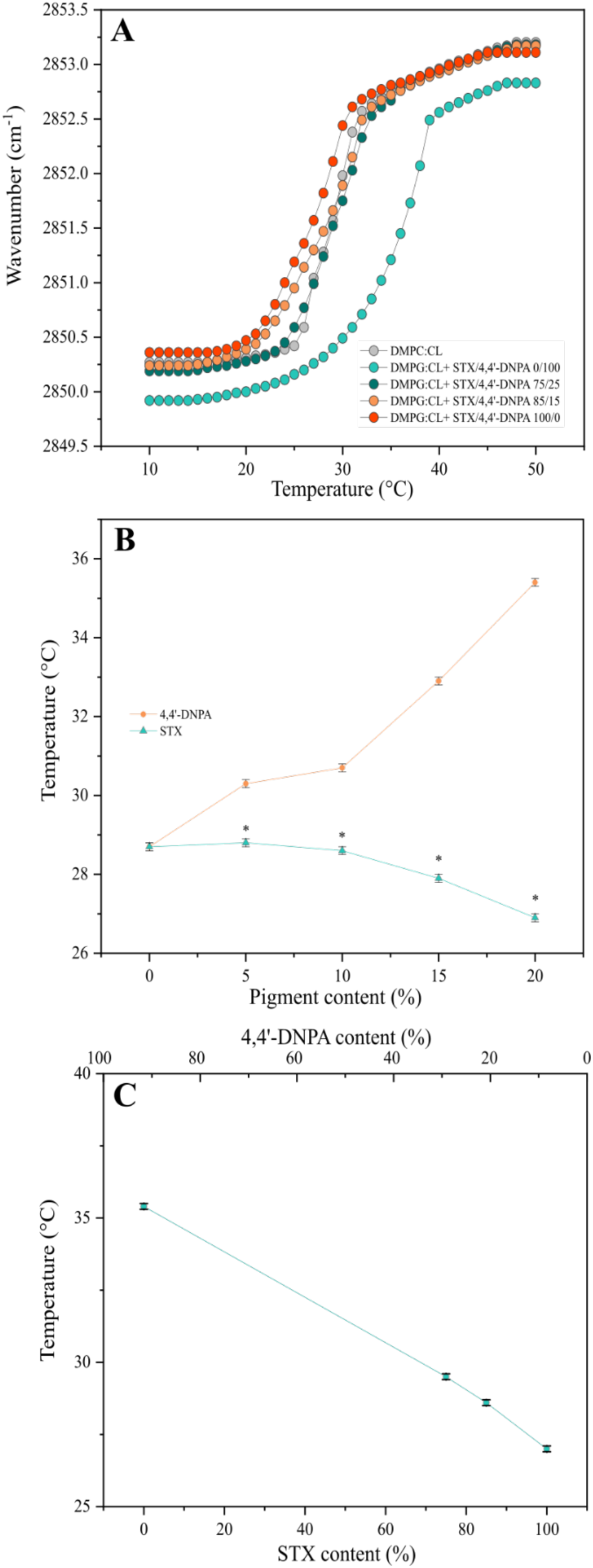
**(A)** Temperature dependence of the peak position of the νCH₂ symmetric stretching band of acyl-chain methylene groups in DMPG:CL models containing STX/4,4′-DNPA mixtures at different ratios. **(B)** Comparison of the individual effects of STX and 4,4′-DNPA on DMPG:CL model membranes. Data marked with an asterisk **(C)** correspond to values previously reported (Múnera-Jaramillo *et al.*, 2024). **(D)** Dependence of the main phase transition temperature on the STX/4,4′-DNPA ratio in DMPG:CL membranes.

### 3.3. Fluorescence spectroscopy

#### 3.3.1. Generalized polarization measurements

The influence of the biosynthetic precursor on the hydrophobic/hydrophilic interface of *S. aureus* membrane lipid models was evaluated using Laurdan generalized polarization. Figure 5A shows the Laurdan GP values as a function of temperature for phospholipid systems (DMPG:CL, 80:20) containing 0, 5, 10, 15, and 20 molar% of 4,4’-DNPA. The L_β_-to-L_α_ phase transition was marked by a pronounced decrease in polarization values, shifting from approximately 0.63–0.60 in the ordered state to 0.118–(−)0.03 in the disordered state.

**Fig. 5.**
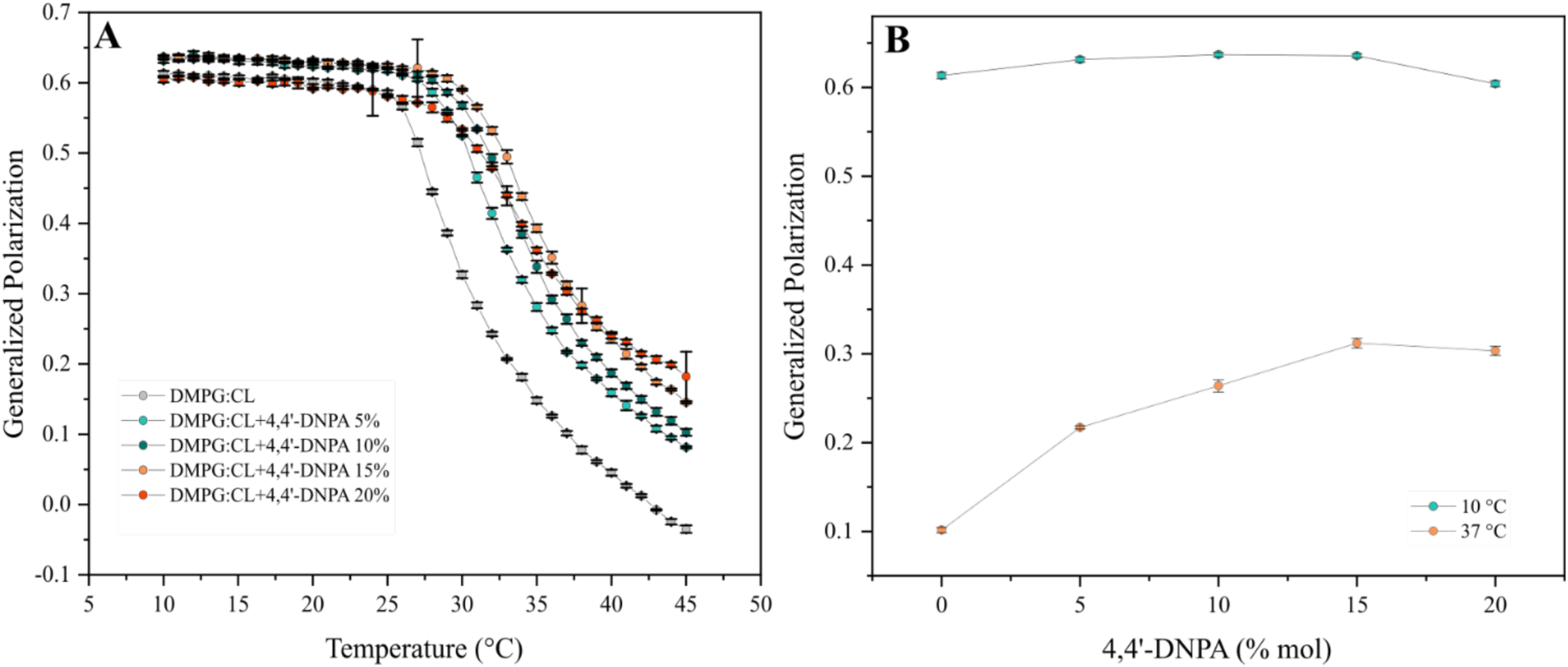
Effect of 4,4’-DNPA in the headgroup spacing of a representative model system of *S. aureus* lipid bilayer. **(A)** Laurdan generalized polarization plotted as a function of temperature for 80:20 DMPG:CL mixtures with increasing amounts of 4,4’-DNPA. **(B)** Generalized polarization variations below (10 °C) and above (37 °C) of the T_m_ for 80:20 DMPG/CL mixtures with increasing amounts of 4,4’-DNPA.

This behavior is characteristic of changes in interfacial hydration during the phase transition, where at lower temperatures reduced hydration is expected, due to increased packing in the solid-ordered phase, while higher temperatures favor increased hydration due to increased headgroup spacing in the liquid-disordered phase [21, 36]. These two-phase behaviors are separated by a cooperative melting event that appears as an inflection in the curve. Figure 5B compares the GP values for lipid systems at 10 °C and at 37 °C, for the different concentrations of the biosynthetic precursor. The results show that below the phase transition temperature, GP values remained largely unchanged among the tested systems, suggesting that the pigment did not substantially alter the hydration in the solid-ordered phase. In contrast, above T_m_, a clear increase in GP values was observed with increasing acid concentration, with the most pronounced effect at 15% acid at 37 °C. The increase in GP values at higher temperatures suggests reduced polarity at the headgroup region, which can be interpreted as decreased interfacial hydration and increased lipid packing at the bilayer interface [21, 37].

#### 3.3.2. DPH anisotropy measurements

The dynamics of the hydrophobic core of the representative models were evaluated through DPH anisotropy [38, 39]. Figure 6A shows the anisotropy values as a function of temperature for 80:20 DMPG:CL systems containing 0, 5, 10, 15, and 20 molar% 4,4’-DNPA. The phase transition from a solid-ordered phase (L_β_) to a liquid-disordered phase (L_α_) was characterized by a notable decrease in DPH anisotropy, with values ranging from 0.31–0.27 to 0.06–0.03, respectively. These results indicate that increasing temperature promotes the rotational dynamic of the probe, reflecting increased mobility and disorder of the lipid acyl chain [39–41].

**Fig. 6.**
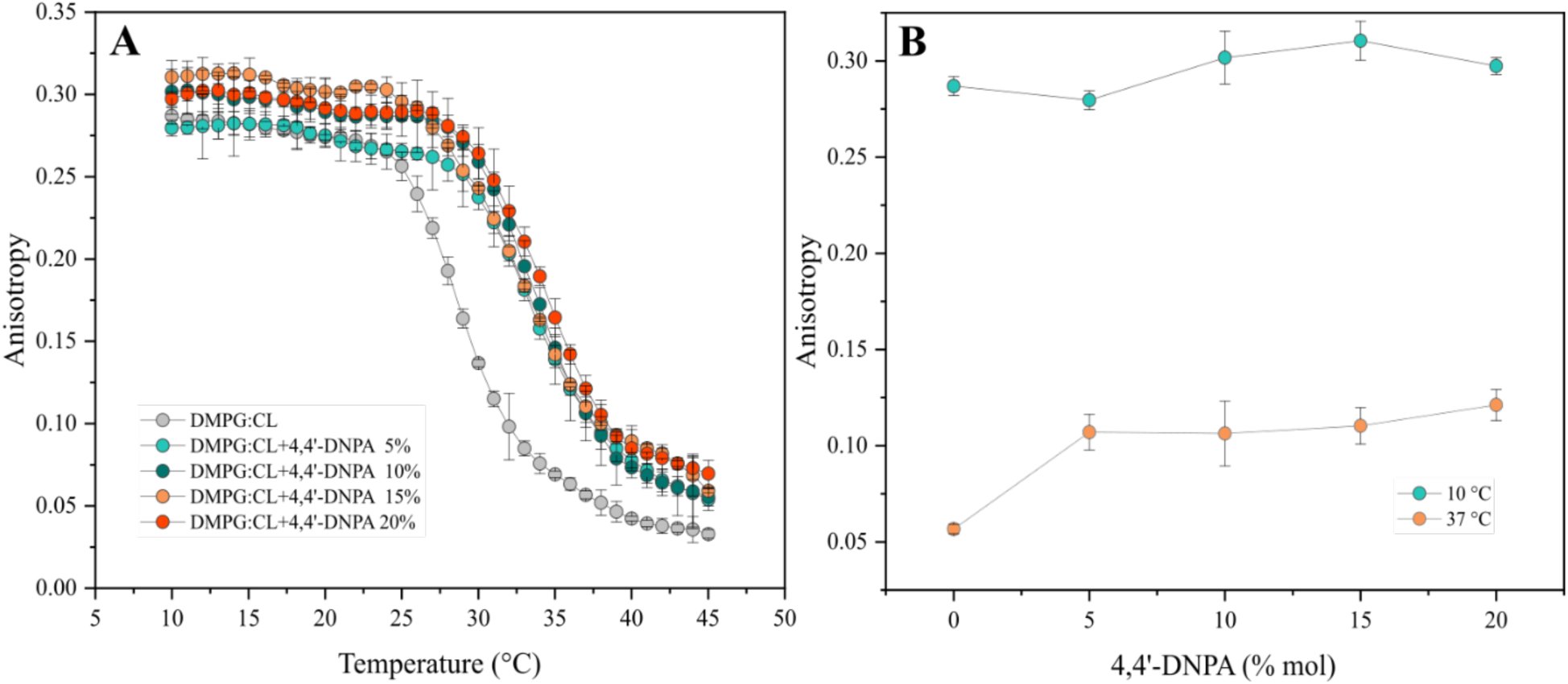
Effect of 4,4’-DNPA in the core dynamic of a representative model system of *S. aureus* lipid bilayer. **(A)** DPH anisotropy measurements as a function of temperature for 80:20 DMPG/CL mixtures with increasing amounts of 4,4’-DNPA. **(B)** Anisotropy changes below (10 °C) and above (37 °C) of the T_m_ for 80:20 DMPG/CL mixtures with increasing amounts of 4,4’-DNPA.

Figure 6B shows the DPH anisotropy values of lipid systems at 10 °C and 37 °C as a function of the concentration of the biosynthetic precursor. Below the phase transition temperature, the systems containing 10% and 15% 4,4′-DNPA exhibited the highest *r* values, indicating a greater restriction of acyl chain motion and increased packing of the lipid molecules [21]. This suggests that, even in the solid-ordered phase, 4,4′-DNPA reinforces bilayer rigidity [21, 28]. At temperatures above T_m_, although the overall thermotropic profile of the systems remained relatively similar for the different acid concentrations, the pigmented models consistently maintained higher anisotropy values than the control. This observation implies that 4,4′-DNPA promotes increased acyl chain order and reduced mobility in the membrane with the strongest effect observed in the liquid-disordered phase. These findings highlight the capacity of 4,4′-DNPA to modulate membrane physical properties over a broad temperature range.

## 4. Discussion

*S. aureus* carotenoids are not only responsible for the characteristic golden pigmentation of this bacterium but also contribute to its pathogenicity [10, 12]. In addition to their antioxidant activity [11, 42], these molecules have been implicated in modulating the biophysical properties of membranes, such as fluidity, packing, and rigidity, thereby enhancing the structural stability and functional integrity of the membrane under stress conditions [15, 17]. In a previous study, the role of the bacterial carotenoid STX as a biophysical regulator was evaluated; STX is known to be the major product of the carotenoid biosynthetic pathway in *S. aureus* [7, 14]. In addition to the purification and characterization of STX by LC-MS, the presence of the persistent biosynthetic precursor 4,4’-DNPA was also evidenced based on chromatographic, UV–Vis, and mass spectrometric data. In this study, the effect of 4,4’-DNPA on the biophysical properties of synthetic lipid models of *S. aureus* cell membranes was investigated. To this end, the bacterial pigment was purified from a total carotenoid extract obtained directly from cells using PTLC. The separation between STX and 4,4′-DNPA was achieved based on their differences in polarity and chromatographic behavior [29]; the biosynthetic precursor contains a polar carboxylic group, in contrast to the glycosylated moiety present in STX. These structural differences are consistent with their distinct chromatographic retention and mass spectrometric behavior, supporting their differential interactions within the lipid bilayer. The coexistence of STX, 4,4′-DNPA, and other carotenoid species in the stationary phase of bacterial growth is supported by LC-DAD-APCI-MS/MS profiles and is consistent with previous reports, which indicate that the characteristic pigmentation of *S. aureus* results from a mixture of several chemical species depending on factors such as the growth medium and other environmental conditions [8,17].

On the other hand, the coexistence of both pigmented species in the stationary phase could suggest a biological mechanism by which *S. aureus* regulates the biophysical properties of its membranes by modulating their proportions [43, 44]. Biological membranes are highly dynamic structures whose functionality depends on physical parameters, such as lateral mobility, mechanical elasticity, and lateral organization of lipids, which determine the membrane’s capacity to withstand environmental changes, host proteins, and maintain vital processes, including transport and signaling [45, 46]. These parameters are strongly influenced by external factors, including temperature, which induces changes in the physical behavior of the lipid components in the bilayer. When a lipid system is cooled, the acyl chains in the lipid components enter a more ordered state with a reduced number of rotamers. This leads to decreased headgroup spacing and reduced lateral mobility. When the lipid system increases in temperature, the acyl chains increase the number of rotamers and the lipid bilayers become more fluid [47, 48]. This event tends to be cooperative, and the temperature at which this phase change occurs is known as the phase transition temperature. The phase transition temperature marks the main lipid phase transition in which the lipid bilayer transitions from a highly ordered state (P_β’_) to a disordered and mainly fluid state ( L_α_) [48, 49].

The phase transition temperature was determined by infrared spectroscopy. The FT-IR results showed that 4,4’-DNPA significantly influences the thermotropic properties of the lipid models evaluated. The most pronounced effect occurs in the pure DMPG lipid system; where a notable increase in the phase transition temperature is observed in the presence of increasing amounts of 4,4′-DNPA. Additionally, the inclusion of high concentrations of the biosynthetic precursor leads to a decrease in the cooperativity of the phase transition. This behavior suggests that 4,4’-DNPA may induce phase separation in the model systems, resulting in a more gradual and less cooperative phase transition at the molecular level [27, 50]. On the other hand, the effect of 4,4’-DNPA on CL bilayers is like that observed in pure DMPG bilayers, although with a markedly lower magnitude in the induced change in the phase transition temperature. This observation is consistent with previous reports indicating that CL, which possesses four hydrophobic acyl chains rather than the two found in typical phospholipids like DMPG, promotes stronger intermolecular interactions, resulting in a more rigid membrane with enhanced mechanical stability [35, 51].

The results obtained for representative model mixtures of *S. aureus* cell membranes showed a progressive increase in rigidity as a function of bacterial pigment concentration, consistent with the distinct physicochemical properties of the carotenoid species putatively identified by LC-DAD-APCI-MS/MS. This behavior differs from previous studies, in which the incorporation of either a mixture of carotenoids or purified STX in lipid bilayers depressed the main transition temperature (Tₘ) [25, 28], indicating greater fluidity. In contrast, 4,4’-DNPA appears to induce the opposite effect, increasing the rigidity of the system.

The phase transition behavior observed in multicomponent systems is of particular significance. According to the results, models containing both STX and 4,4’-DNPA show a phase transition profile comparable to that of the control system, suggesting that the accumulation of acid during the stationary phase may act as a regulatory mechanism to counteract the fluidizing effect of STX. The above is relevant in biological terms, as cell membranes must maintain an intermediate physical state to ensure their functionality: sufficiently fluid to allow for dynamism and molecular reorganization, but stable enough to preserve structural integrity [45, 46].

The observed differences in the effects of the two carotenoids on the lipid model systems can be rationalized in terms of their structural characteristics. A comparison of the chemical structures of 4,4′-DNPA and STX evidence that both molecules share the characteristic diaponeurosporenoic acid chain. However, unlike STX, 4,4′-DNPA lacks both the glucose ring and the additional aliphatic chain attached to it. The absence of the bulky glycoside moiety alters the interactions between the pigment and the phospholipids in the polar region of the bilayer, and consequently, the ordering within the hydrophobic core. The smaller polar group of 4,4′-DNPA, compared to the bulky glucose in STX, facilitates closer and more effective interactions with the lipid headgroups, promoting tighter packing. This results in increased molecular organization of the membrane, yielding a more ordered lipid bilayer.

Changes in lipid and pigment content influence the hydration levels at the hydrophilic/hydrophobic interface, as well as the degree of ordering and dynamics of the acyl chains within the hydrophobic core [52, 53]. These effects can be probed using environment-sensitive fluorescent dyes such as Laurdan and DPH, which are widely employed to study membrane biophysics [41]. The fluorescent moiety of Laurdan locates at the interfacial region of the bilayer, at the boundary between the hydrophobic core and the hydrophilic headgroup region, with its fluorophore positioned near the glycerol backbone and upper acyl chain region, where it senses polarity changes due to water penetration and headgroup spacing [54, 55]. Consequently, Laurdan generalized polarization (GP) values decrease as interfacial hydration increases, reflecting the transition from an ordered, dehydrated gel phase to a disordered, hydrated liquid-crystalline phase [21, 36]. In parallel, DPH inserts deep into the hydrophobic core of the bilayer, locating along the acyl chains, and reports on the rotational dynamics of this fluorescent probe [41]. High DPH anisotropy values correspond to tightly packed, ordered chains in the gel phase that curtail rotational mobility of DPH, whereas low values indicate increased chain dynamics and disorder in the fluid phase, with high mobility of DPH [21, 39].

The interface hydration and the degree of order in the hydrophobic core were tested using Laurdan and DPH probes, respectively. The Laurdan GP results showed a decrease in interface hydration as pigment concentration increased, particularly at high temperatures.

This effect suggests a strong interaction between the polar carboxylic group of 4,4′-DNPA and the polar heads of the phospholipids, which can displace water molecules from the interface, where the carboxylic group would provide a negative charge to 4,4′-DNPA [37, 56]. In this case, 4,4′-DNPA, although charged, lacks a bulky headgroup, therefore inducing lipid condensation and the headgroup region. This behavior is consistent with reports indicating that small polar groups tend to form hydrogen bonds with the head groups of phospholipids, leading to local surface dehydration [57, 58]. In the case of DPH anisotropy measurements, 4,4′-DNPA induces notable changes in the bilayer core. These findings indicate that, at higher concentrations, 4,4’-DNPA significantly restricts the dynamics of the acyl chains, increasing their order and reducing their molecular motion [41]. This behavior is consistent with the rigid and linear nature of the triterpenoid chain, which intercalates between lipid chains, restricting their lateral and conformational movements [18, 28]. Similar behavior has been reported for other carotenoids in model membranes, where they increase chain order and reduce bilayer fluidity [19, 21, 59]. The observed reduction in interfacial hydration, together with the increased order of the hydrophobic core as a function of pigment concentrations, supports the idea that 4,4′-DNPA modulates the physicochemical properties of the bilayer, stabilizing both the hydrophobic and the polar regions. Notably, these effects are more pronounced at elevated temperatures, which can be attributed to the bilayer adopting a more fluid and disordered state [60], thereby becoming more susceptible to carotenoid-induced reorganization.

The results suggest that the coexistence of 4,4′-DNPA and STX in the stationary phase may be a homeoviscous regulatory mechanism of *S. aureus* to maintain the optimal physicochemical properties of their lipid bilayers [61, 62]. Homeoviscous adaptation is a well-known strategy in microorganisms to preserve membrane functionality by adjusting lipid composition and bilayer organization in response to environmental changes, such as temperature changes [61, 62]. In this context, where the membrane requires greater order and rigidity, 4,4′ DNPA, with its small and highly polar carboxylic group, appears to integrate into the lipid bilayer, promoting more efficient packing of lipid chains and decreasing hydration at the interface. On the other hand, when the membrane is too rigid, the conversion of 4,4′ DNPA into staphyloxanthin, through the action of the enzyme CrtQ (Fig. 1) [7], produces a molecule with a more bulky polar group that could introduce greater disorder and modulate rigidity at the interface. This balance between the two carotenoids represents a regulatory strategy that enables *S. aureus* to adjust its chemical composition, thereby modulating the fluidity and organization of its membrane in response to environmental changes. Thus, the enzymatic control of carotenoid biosynthesis results in the structural regulation of the membrane, ensuring its functionality and stability under conditions of external stress [63].

## Conclusions

In this study, we evaluated the effect of the carotenoid biosynthetic precursor 4,4′-DNPA on the thermotropic and biophysical properties of representative *S. aureus* membrane models. The results demonstrate that 4,4′-DNPA modulates the physical state of these systems increasing acyl chain order, reducing interfacial hydration, and enhancing membrane rigidity, as evidenced by elevated phase transition temperatures and reduced core dynamics. In contrast, the main carotenoid, STX, decreases the transition temperature and promotes membrane fluidity. The combined presence of both pigments in multicomponent systems reveals a dynamic equilibrium, in which their relative proportions regulate membrane organization, suggesting a homeoviscous adaptation mechanism that maintains an optimal balance between fluidity and stability. These findings highlight that carotenoid intermediates act not only as biosynthetic precursors but also as active modulators of membrane structure and function, providing evidence that carotenoid production serves as a biophysical regulatory mechanism in *S. aureus* membranes. This capacity to adjust membrane properties likely contributes to the ability of *S. aureus* to survive and adapt under changing environmental conditions.

## Abbreviations

S. aureus: Staphylococcus aureus
STX: staphyloxanthin
AMPs: antimicrobial peptides
FT-IR: Fourier-transformed infrared spectroscopy
DSC: differential scanning calorimetry
4,4’-DNPA: 4,4’-diaponeurosporenoic acid
DMPG: 1,2-dimyristoyl-sn-glycero-3-phosphoglycerol sodium salt
CL: 1’,3’-bis [1,2-dimyristoleoyl-sn-glycero-3-phospho]-glycerol sodium salt
HEPES: 4-(2-hydroxyethyl)-1-piperazineethanesulphonic acid
HPLC: high-performance liquid chromatography
PLE: pressurized liquid extraction
NaCl: sodium chloride
BHT: butylated hydroxytoluene
EDTA: ethylenediaminetetraacetic acid
LB: Luria-Bertani
LAURDAN: 6-Dodecanoyl-2-Dimethylaminonaphthalene
DPH: 1,6-Diphenyl-1,3,5-hexatriene
PFTE: poly-tetrafluoroethylene
PTLC: preparative thin liquid chromatography
LC-MS: liquid chromatography-mass spectrometry
DAD: diode-array detection
MS: mass spectrometry
APCI: atmospheric pressure chemical ionization
SLBs: supported lipid bilayers
MCT: Mercury-Cadmium-Telluride
T_m_: transition temperature
MLVs: multi-lamellar vesicles
GP: generalized polarization
r: anisotropy
L_β_: solid-ordered phase
L_α_: liquid-disordered phase.

## Conflict of Interest

The authors declare that they have no known competing financial interests or personal relationships that could have appeared to influence the work reported in this paper.

## Acknowledgements

The research was funded by MinCiencias through the Grant Program Grant Program (Cod. 120480763040, RC 846-2018). Additional support came from an internal grant from the Faculty of Sciences at Universidad de los Andes (INV-2025-213-3435). We would like to thank the Faculty of Sciences for granting the Academic Semester for Research (STAI semester 2026-1), which provided valuable time for completing this manuscript.

## Notes

### Competing Interest Statement

The authors have declared no competing interest.

