## Supplemental Figs 1S and 2S for "Membrane structural properties in *Staphylococcus aureus* are tuned by the carotenoid 4,4′-diaponeurosporenoic acid"

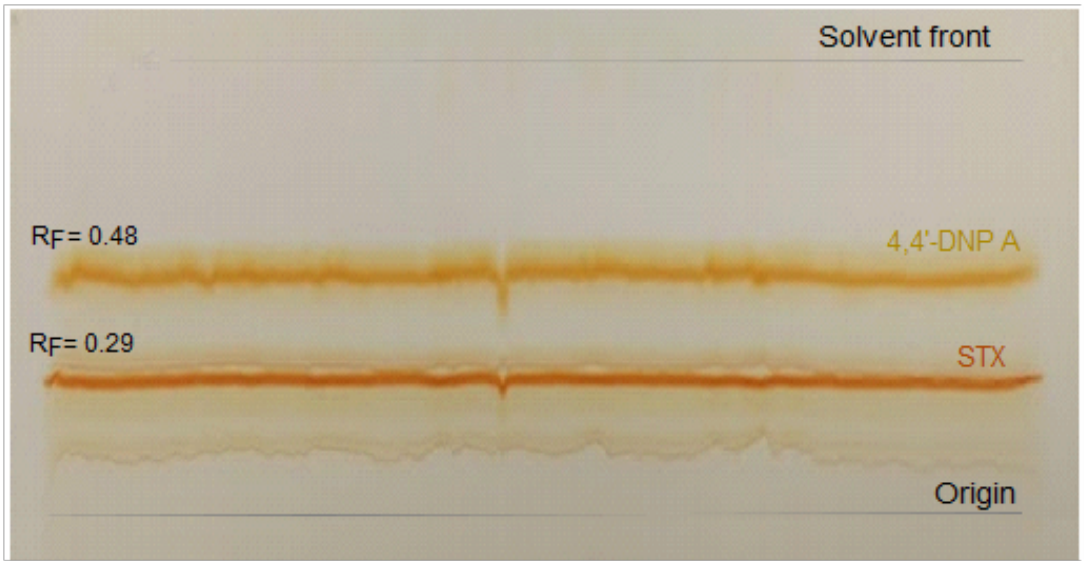


**Fig 1S.** Plate of bacterial carotenoids separation from *S. aureus* cells by PTLC. The yellow band is attributed to 4,4'-DNPA based on its chromatographic behavior and subsequent LC-MS analysis, while the orange band corresponds to staphyloxanthin, consistent with its characteristic pigmentation.

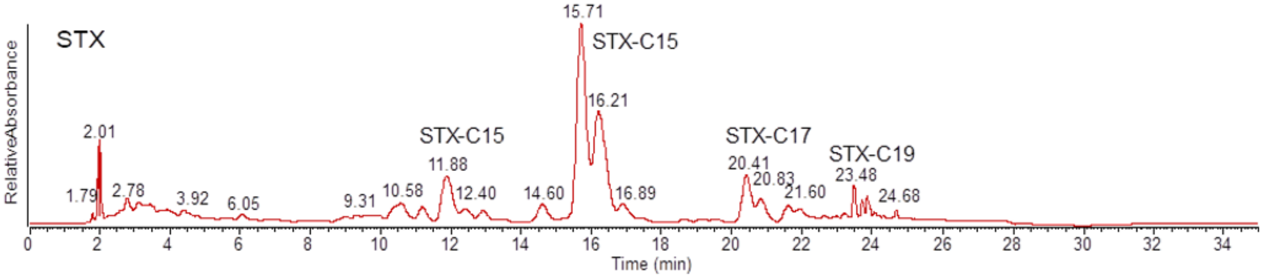


**Fig 2S.** LC-DAD-APCI-MS/MS analysis of purified STX. The detected peaks correspond to STX homologs (e.g., STX-C17 and STX-C19), based on their chromatographic retention and APCI-MS/MS fragmentation patterns, consistent with previous reports. .
